# A viral vector model for circuit-specific synucleinopathy

**DOI:** 10.1101/2023.08.28.555129

**Authors:** Alexia Lantheaume, Nina Schöneberg, Silvia Rodriguez-Rozada, Dennis Doll, Michael Schellenberger, Konstantin Kobel, Kilian Katzenberger, Jérémy Signoret-Genest, Angela Isabel Tissone, Chi Wang Ip, Maria Soledad Esposito, Philip Tovote

## Abstract

In Parkinson’s disease (PD), pathomechanisms such as aberrant network dysfunctions can be elucidated by conducting multiscale explorations in animal models. However, the lack of specificity in the existing models limits a restricted targeting of individual network elements and characterization of PD as a “circuitopathy”. We therefore developed a cell-type specific viral vector (AAV2/9-Cre^ON^-A53T-αSyn) mouse model that allows to induce synucleinopathy within individual circuit elements *in vivo*. When specifically targeted to dopaminergic (DA) neurons of the substantia nigra pars compacta (SNc), our approach recapitulates the main hallmarks of the disease, namely Lewy-body-like aggregation, progressive cellular and nigrostriatal projections loss, together with locomotor impairment. Our strategy is supported by new state-of-the-art analytical approaches for cell quantification and behavior characterization. Altogether, we provide a novel model of synucleinopathy, which offers new opportunities to study the contribution of individual network elements to disease pathomechanisms.

## INTRODUCTION

As the main treatment approach of synucleinopathies such as Parkinson’s disease (PD), current pharmacotherapies target the dopamine metabolism to functionally correct deficiency of this neuromodulator^1^. However, this option still suffers from many caveats (e.g. treatment resistance, dystonia)^2,3^ and preventive treatments are currently missing due to the lack of detailed knowledge of the pathomechanisms and disease etiology. Because of the resolution animal models offer for assessing mechanistic details, they are a substantial part of PD research^4–7^.

Historically, PD was exclusively attributed to dopaminergic cell death within the substantia nigra pars compacta (SNc). From this “lesion hypothesis”, the first exploratory studies using neurotoxin-based animal models originated^8^. Although these mouse models phenocopy behavioral motor effects of PD, given their rapid mode of action and induction of effects^9^ they fall short in capturing the progressive neurodegenerative pathomechanisms underlying PD.

At the molecular level, the aggregation of α-synuclein (αSyn) is a major component of PD pathophysiology and is thought to promote cell death^10,11^. αSyn is consistently found overexpressed in the SNc of PD patients, with the A53T mutation presenting the prime example of hereditary synucleinopathy resulting in PD^12^. Notably, genome-wide association studies indicated a close association between genetic variability of the αSyn locus and PD risk^13^. This finding motivated the development of numerous animal models for PD, using αSyn overexpression and/or mutation via different promoters and transgenes^6^. The emergence of genetic models provided new insights into the development of cellular dysfunctions, concomitant to motor and non-motor symptoms^14^. More recently, viral vector-mediated expression of pathogenic A53T-mutated human αSyn (pαSyn) was the first approach to produce a robust αSyn-driven degeneration of SNc neurons in mammals, alongside with strong behavioral defects^15^. Another major advantage was a quick and reliable time frame of development of cellular and behavioral dysfunctions as compared to other transgenic approaches.

Nevertheless, most of the research conducted to date studied PD pathomechanisms either at the molecular level, in cellular models, or at the level of entire brain regions^16^. Recent studies have started to address the neuronal circuit level as a main unit of brain function, and thus crucial to complement our understanding of the disease. They reveal how not only individual brain regions, but brain-wide networks are impacted^16^, characterizing PD as a “circuitopathy”^17^. This is reflected, for example, by differences in the sensitivity among dopaminergic neurons to αSyn overexpression^18^. Those emerging concepts highlight the importance of cell-type heterogeneity within brain areas, as well as the synaptic connections from major cellular elements affected in PD with other neuronal populations. Hence, models that allow cell-type-specific induction of PD pathomechanisms are required to dissect the contribution of individual circuit elements to PD symptoms^17^. Consequently, we generated a conditional adeno-associated viral vector (AAV) that allows expression of pαSyn in selected neuronal subtypes using the Cre-lox site-specific recombination system^19,20^. We specifically targeted dopaminergic (DA) neurons of the SNc through local intracranial injection of the pαSyn AAV in DAT-Cre transgenic mice.

The variety of symptoms highlights PD as a progressive systems disease. Besides the motor symptoms, a diverse array of non-motor symptoms exists in PD, with some of them serving as prodromal manifestations preceding the onset of the disease^22,23^. As a major component, cardiac autonomic dysfunction often worsens with disease progression^22^. It has been shown that heart rate (HR) parameters and variability are decreased in PD patients^24,25^. Although cardiac parameters as potential biomarkers of synucleinopathy may help understanding disease onset and developing early neuroprotective options^26^, they have rarely been assessed in animal models of PD. Thereupon, we measured the evolution of HR variables in freely moving mice in the time course of pαSyn expression in the SNc.

We demonstrate that this approach enables cell-type specific targeting and induces major cellular hallmarks of synucleinopathy. Indeed, it initiates αSyn phosphorylation and cell death within the SNc, loss of nigrostriatal projections, alongside with behavioral impairments. The dysfunctions occurred within a similar timeline to that seen when using unspecific viral approaches^15^. For quantification of the neurodegenerative and behavioral effects, we used customized state-of-the-art, machine-learning image and video analysis to avoid observer bias and ensure a robust and highly reliable validation of our viral tool^27^. Taken together, our results present a novel Cre-driven genetic model of synucleinopathy constituting a valuable tool for exploring the network effects of αSyn aggregation, thereby facilitating our understanding of PD as a circuitopathy.

## RESULTS

### Inducing cell-type specific synucleinopathy: approach and analysis

A Cre recombinase-conditional AAV was designed to target specific neuronal subtypes selected by local administration in Cre-mouse driver lines. Briefly, a human pαSyn (A53T mutated) complementary DNA sequence fused with a Myc-tag was placed in reversed orientation between double inverted loxP sites (*Fig. 1a*). The construct was encapsulated into an AAV2/9 (AAV2/9-Cre^ON^-pαSyn). To assess DA-SNc neuronal loss upon pαSyn expression, we injected the AAV unilaterally into the SNc of mice expressing Cre recombinase under the promoter of the dopamine transporter gene (DAT-Cre transgenic mouse line). Blue-fluorescent beads were co-injected to verify correct targeting within the SNc. At 8 and 10 weeks after injection mice were sacrificed and DA-SNc cells were identified via tyrosine hydroxylase (TH) immunostaining (*Fig. 1b, Fig. S1*).

**Fig. 1.**
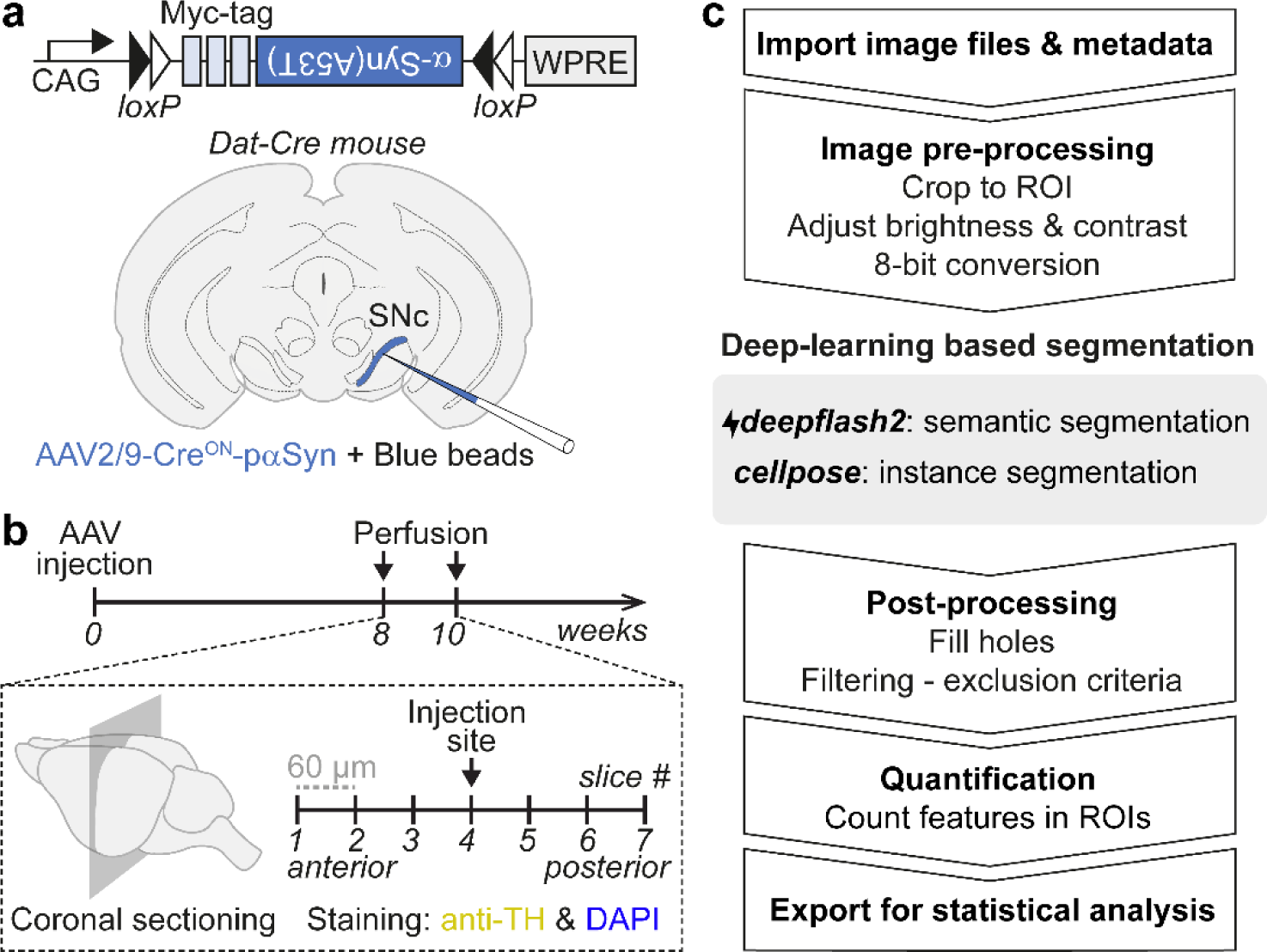
Development of viral and image analysis tools. **(a)** Cre-dependent viral strategy to achieve specific expression of A53T-αSyn in dopaminergic neurons of the SNc. Mice injected with an empty vector served as a control group. **(b)** Experimental timeline. Blue beads were used to identify the injection site. **(c)** Analysis pipeline for quantification of TH+ cells. The deep-learning based segmentation step uses the previously developed algorithms *deepflash2* and *cellpose*. See Fig. S2 for a detailed step-by-step explanation of *findmycells*.

While TH+ cell labelling and quantification has been commonly used as a proxy of pαSyn-induced cell death, manual annotation of fluorescent features can not only be time-consuming but also subjective^28^. Novel machine-learning based image analysis tools help overcome this issue but often require programming skills, thereby hampering their use by non-experts. Here, we developed a new analysis pipeline, *findmycells*, which combines modern deep-learning algorithms with several image preprocessing options in an easy-to-use graphical user interface (GUI), thus allowing comprehensive, unbiased analysis of large image datasets in a standardized and effective way (see *Fig. S2* for detailed description).

We then took advantage of some key features of *findmycells* to quantify TH+ cells after pαSyn expression (*Fig. 1c*). Briefly, a uniform dataset was created by adjusting brightness and contrast across all 8-bit images, which was then used for deep-learning based segmentation. After training a *deepflash2*^27^ expert-consensus ensemble on a subset of TH staining images, TH+ cells were reliably identified by semantic segmentation. Individual instances predicted by *cellpose*^29^ further improved segmentation accuracy.

### Expression of pαSyn in dopaminergic SNc cells leads to cell death and loss of nigrostriatal projections

Two cardinal features characterize PD histopathology: the progressive loss of DA-SNc neurons, and the presence of Lewy bodies which consist of intracellular protein aggregates enriched in phosphorylated αSyn^11,14^. To assess whether the expression of the mutant (A53T)-αSyn can recapitulate both features, we first quantified the density of TH+ neurons in SNc using the quantification results by *findmycells* from the respective images *(Fig. 2a, b)*. We found that conditional expression of pαSyn for 10 weeks leads to a significant reduction in the density of TH+ SNc neurons in comparison to DAT-Cre mice injected with an AAV1/2-empty vector (EV, *p= 0.03, Fig. 2c)*. To quantify potential DA loss differences, brain slices from mice injected with pαSyn were grouped into 3 different categories based on their localization to the injection site: anterior (2 slices), medial (3 slices), posterior (2 slices). When comparing ipsilateral (pαSyn) to contralateral non-injected side, the anterior part of the SNc registered the highest TH+ cell loss (*p= 0.01*), followed by the medial part of the SNc (*p= 0.04*), while no significant DA-cell loss was observed in the posterior sections of the SNc (*p=0.32, Fig. S3a*). These results are consistent with the fact that cell numbers vary within SNc and are lower in the posterior parts^30^. We next performed double immunostaining against the Myc-tag to identify exogenous αSyn expression, and against phosphorylated human αSyn at serine-129 residue (pS129), as a Lewy-body marker. We observed colocalization between the Myc-tag and pS129, supporting the development of human Lewy-body-like pathology^11,14^ (*Fig. 2d).* In line with Ip et al.^14^, the strongest signal could be visualized in the cytoplasm of the cells.

**Fig. 2.**
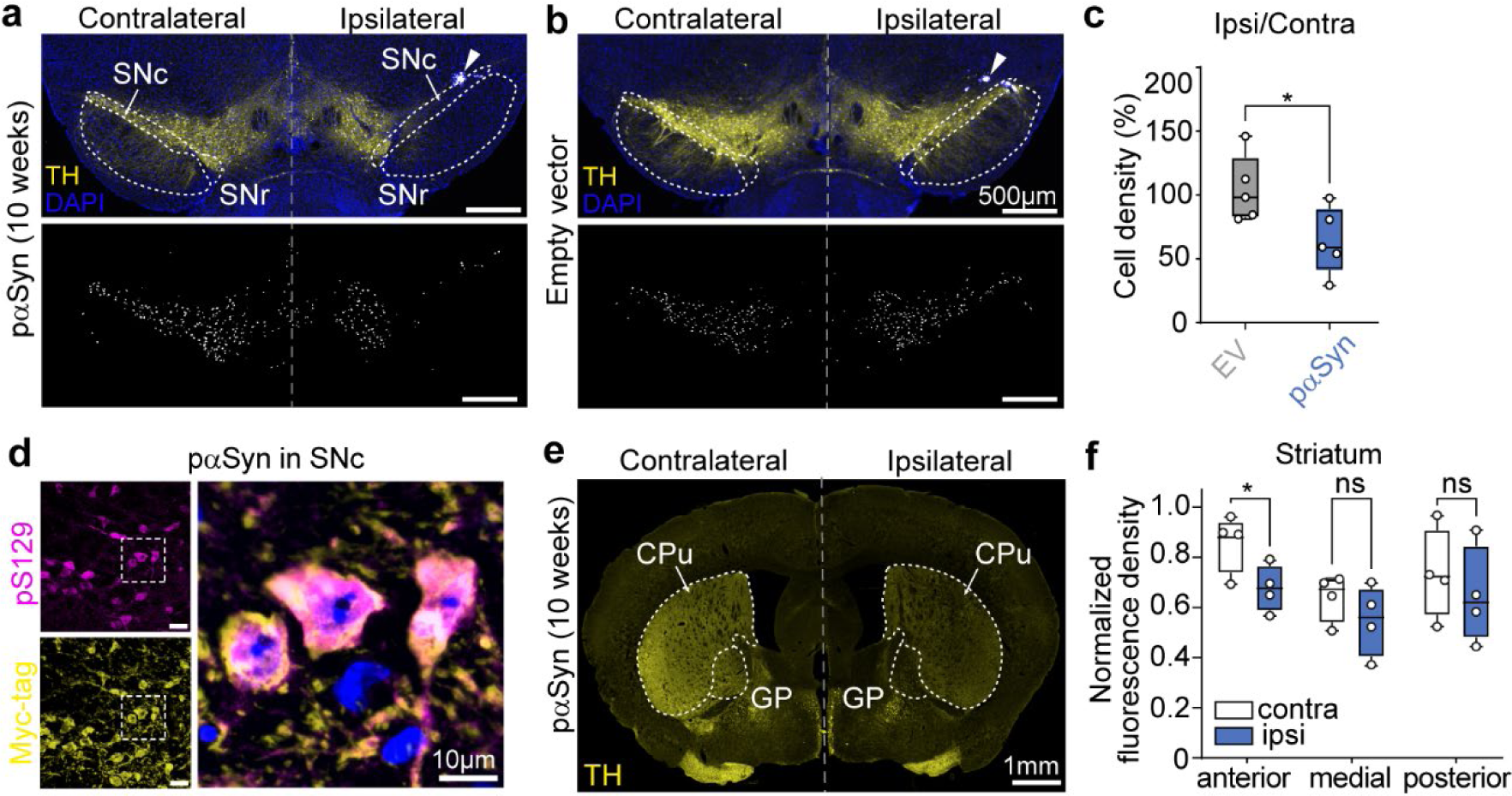
Neurodegeneration of dopaminergic SNc cells upon cell-specific expression of pαSyn. **(a)** Representative maximum-intensity projection image of immunostainings showing TH expression (yellow) in the SNc of a DAT-Cre mouse 10 weeks after unilateral injection of Cre-dependent pαSyn (top). Blue beads were co-injected to identify the injection site (white triangle). White dotted lines delineate SNc and substantia nigra pars reticulata (SNr) areas according to the Allen Mouse brain atlas. Bottom: corresponding mask of TH+ cells identified by *findmycells.* **(b)** Same as (a) but for mice injected with the control empty vector (EV). **(c)** Percentage of ipsilateral/contralateral cell density for pαSyn and EV groups 10 weeks post-injection. For each mouse, cell density was quantified in seven 30 µm-thick brain sections and averaged. Dots correspond to mean values per animal. Box plots show mean ± min/max values, n=5 mice for both pαSyn and EV group. Mann-Whitney test. **(d)** Maximum-intensity projection images of a confocal z-stack showing expression of pαSyn (Myc-tag, yellow) and the Lewy-body marker pS129 (magenta) in SNc neurons, scale bar 25 µm (left). Right: magnified view showing colocalization of Myc and pS129. DAPI staining is shown in blue. **(e)** Example image showing TH expression in striatal-projecting SNc axons of a DAT-Cre mouse 10 weeks after unilateral injection of pαSyn in the SNc. Caudate Putamen (Cpu) and Globus pallidum (GP). **(f)** Quantification of TH+ fluorescence density in the ipsi- and contralateral striatum (Cpu) along the anterior-posterior axis. For each mouse, mean fluorescence density was quantified in 6 striatal sections (2 anterior, 2 medial, 2 posterior) and normalized to the highest value. Box plots show mean ± min/max values, n=5 mice, two-way ANOVA with Sidak’s multiple comparisons test (*p<0.05).

To understand how fast the progression of synucleinopathy leads to cell death, we also quantified TH+ neurons 8 weeks after injection of pαSyn. Although at this time point, there was a marked tendency towards lower TH+ cell density as compared to the EV control group, the difference was not significant (*p=0.052, Fig. S3b)*, indicating a lesser advance in the degree of neurodegeneration at a shorter time point. In addition to the pαSyn condition, we tested another cell-type specific vector delivering a wild type (non-mutated) human αSyn (AAV2/9-Cre^ON^-αSyn). After 10 weeks, the cell loss reflected a clear tendency without reaching significance (*p=0.08, Fig. S3b*) and thus constituted an intermediary phenotype when compared to the A53T mutant viral vector and EV. These results are in line with the hypothesis that overexpression of the non-mutated αSyn also leads to cell loss but at longer timescales^31^.

To rule out the possibility that pαSyn overexpression might lead to reduced TH+ expression levels rather than neuronal loss, we performed another series of experiments in which we co-injected in the SNc of DAT-Cre mice AAV2/9-Cre^ON^-pαSyn and another viral vector conditionally expressing GFP (AAV2/9-Cre^ON^-EGFP-KASH) on one side (*Fig. S4*). As a control, we co-injected into the contralateral SNc a mixture of AAV2/9-Cre^ON^-EGFP-KASH and AAV2/9-Cre^ON^-H2B-Tomato. When we quantified the amount of GFP+ neurons on both sides we found a significant reduction of labeled DA-SNc neurons in the side co-injected with pαSyn after 8 weeks of expression, demonstrating univocally that our novel tool can induce the degeneration of DA neurons in the SNc.

Having demonstrated that our virally mediated aggregation of pαSyn leads to local cell loss, we next checked whether cell death of DA cells in the SNc is accompanied by a loss of striatal projections, for which we measured mean fluorescence intensity in the striatum of mice 10 weeks post-injection *(Fig. 2e)*. Fluorescence density of the ipsilateral side was compared with the contralateral side along the anterior-posterior axis (*Fig. 2f).* Injection of pαSyn into the SNc resulted in significant loss of ipsilateral anterior nigrostriatal projections (*Fig. 2f, p=0.01*). Taken together, our findings demonstrate the effectiveness of the Cre-conditional vector in terms of inducing Lewy-body-like pαSyn overexpression, local cell loss, and degeneration of specific axonal projections.

### Specific loss of dopaminergic SNc neurons induces behavioral deficiency

After validating the cell-type specific induction of synucleinopathy by our viral vector, we next assessed its effect on locomotor behavior. To mimic bilateral aggregation of pαSyn in PD, we injected the viral vector into the SNc of both hemispheres (*Fig. 3a*). Mice of the control group were bilaterally injected with a Cre-dependent AAV encoding the fluorescent tag mCherry. Confirming the cell loss evaluation achieved previously, the pαSyn group contained 2.6 times less DA-SNc cells than the control group (respective *average density 0.3 and 0.8, p=0.026, Fig. 3b*), when sacrificed at the end of the behavioral assessment. To determine the progressive effect of pαSyn-induced toxicity on locomotion and motor performance, mice were subjected to standard behavioral assays at several time points (*Fig. 3a, experimental plan*). The rotarod is one of the most common paradigms to assess locomotor skills and has widely been applied in PD models^32^. We trained mice in an accelerating rotarod task one week before surgery and assessed their latency to fall at several time-points after surgery to assess their skill performance. All mice started with a similar average score on week 1 (*Fig. S5a*), and while the control group remained stable throughout the weeks, pαSyn-expressing mice progressively got worse on the task (*Fig. 3c, p=0.037*). When comparing all weeks individually, we observed that already 8 weeks after injection the groups had differing scores (*Fig. S5a*), consistent with the cell loss observed already at week 8 (*Fig. S3b*).

**Fig. 3.**
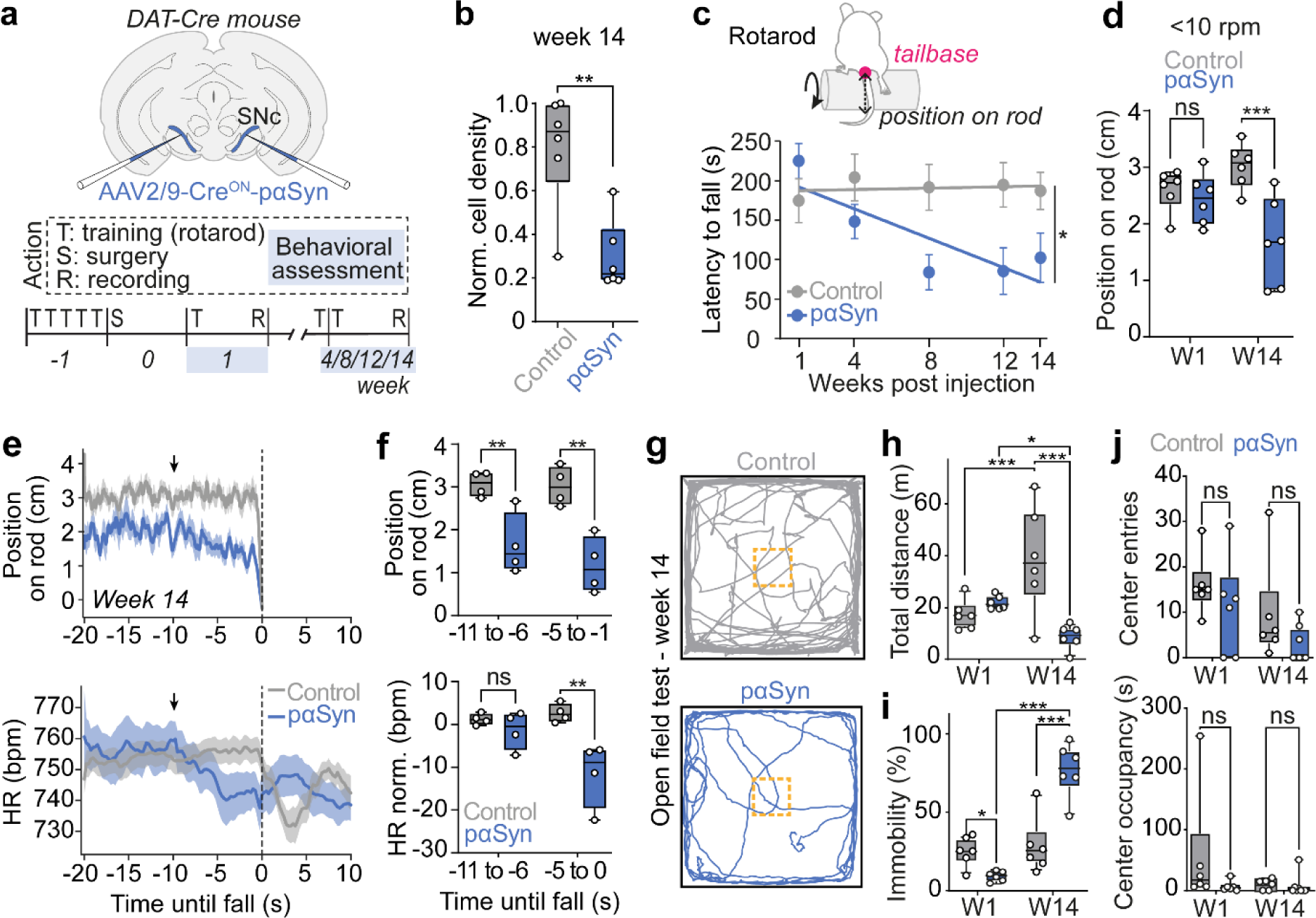
Validation of the cell-specific synucleinopathy model on a behavioral level. **(a)** Experimental design for behavioral assessment. pαSyn was injected bilaterally in DA-SNc neurons. Control mice were injected with a Cre-dependent vector encoding mCherry. Mouse behavior was assessed using the rotarod and OF test at 1, 4, 8, 12 and 14 weeks after virus injection. **(b)** Normalized TH+ cell density in the SNc of control and pαSyn mice following behavioral assessment 14 weeks post-injection. Dots correspond to mean values of 9 brain sections per animal. Box plots show mean ± min/max values, n=6 mice for both pαSyn and control. Mann-Whitney test. **(c)** Top: schematic drawing of a mouse on the rod showing the tracked tail base (magenta dot) to compute mouse position relative to the low extremity of the rod (in cm). Bottom: latency to fall during the rotarod test at successive weeks post-injection. Dots correspond to mean values ± SEM (n=8 and 7 mice for pαSyn and control, respectively). Linear regression slope comparison. **(d)** Mean tail base position during rotarod trials at slow rotation (6-10 rpm) at weeks 1 (W1) 14 (W14). Box plots show mean ± min/max values, n=6 mice for both pαSyn and control. Mixed model with Bonferroni tests. **(e**) Peristimulus time histogram (PSTH) of the position of the mouse on the rod (Top) and heart rate (Bottom) from 20 s before, until 10 s after the fall at W14. Dashed line shows when the fall occurs. **(f)** Comparison of mean position on the rod (Top) and heart rate (Bottom) for two timepoints **(g)** Representative trajectories within the OF test for control (top) and pαSyn (bottom) group at W14. The yellow square represents the center area of the maze. **(h)** Quantification of total distance, **(i)** immobility probability, **(j)** center entries and occupancy during OF at weeks 1 and 14. Box plots show mean ± min/max values, n=6 mice for both pαSyn and control. Mixed model with Bonferroni test (*p<0.05, **p<0.01 and ***<0.0001).

To characterize this locomotor deficiency in detail, we video-recorded the mice from the back of the rod and tracked their body parts during the trials, using a trained DeepLabCut^33,34^ network. Then, using a custom-made analysis pipeline, we extracted the distance between the tail base position relative to the bottom right corner of the rod, reflecting the mouse position on the rod (*Fig. 3c, drawing*). Given that pαSyn mice tend to perform worse at later time points, the results were divided into two speed categories: slow (from 6 to 10 rounds per minute - rpm) and fast rotation (10 to 50 rpm). A progressively lower position on the rod, evidenced by changes in the tail base position, was observed at slow rotation in pαSyn mice between week 1 and 14 and compared to control mice at week 14 (*Fig. 3d, p=0.005)*. Overall, pαSyn struggled to stay on the upper part of the rod creating an unsteady position, which increased the risk to fall, even when the rod turns slowly. Interestingly, the control group showed an opposite development over time, with a significantly higher position on the rod in fast rotation at week 14 compared to week 1 (*Fig. S5b)*. This change can be explained by habituation to the task and motor learning processes that allow for an optimally stable position on the rod. Taken together, the detailed kinematic analysis demonstrates that pαSyn mice develop an unstable body adjustment over the course of the disease.

To elucidate the dynamics of the locomotor impairment as part of an integrated state involving concomitant autonomic adjustments, we analyzed mice HR on the rotarod via chronically implanted subcutaneous electrodes. This is visualized by a peristimulus time histogram (PSTH) for the position on the rod and mean HR (*Fig. 3e*), centered on the fall time (timepoint 0, when the mouse position reaches 0.3 cm below the rod). At week 1, both rod position and HR dynamics are similar between groups (*Fig. S6a, b*). At week 14, healthy mice stay on top of the rod until they fall, whereas pαSyn mice exhibit a generally lower rod position and a slight decrease starting 10 s before they fall (*Fig. 3e, f; top panels*). Interestingly, HR values show a marked downward shift 10 s before the fall, indicating a state switch including locomotor and HR change before the fall, which is absent in the control mice (*Fig. 3e, f; bottom panels, p= 0.002*).

To test for general locomotor impairments during self-paced behavior, as well as a potentially more generalized anxiety-related effect of DA-SNc cell loss upon pαSyn expression, mice were subjected to the open field (OF) test. At 14 weeks of expression, mice from the pαSyn group exhibited shorter traveling distances when compared to the control group (*Fig. 3g, h*). While both groups start with a similar speed at week 1, control mice significantly increased their velocity in week 14 (*p=0.005, Fig. 3h)*, whereas pαSyn demonstrated slower locomotion speed (*p=0.03*), with values slower than in control mice at week 14 (*p=0.001*). Complementing this result, even though control mice initially were more immobile than the pαSyn-expressing mice at week 1 (*p=0.02)*, the trend was inverted at week 14. In this progressed state of pαSyn aggregation and induced neurodegeneration, mice freeze significantly more when compared to controls (*p<0.0001*) or to themselves in week 1 (*p<0.0001, Fig. 3i).* To investigate whether those differences are reflecting that pαSyn mice become more anxious, we checked their behavior in the center of the OF, the most stressful area for rodents. Remarkably, neither the number of entries, nor the occupancy of this area differed across groups and across time (*Fig. 3j)*. Thus, the reduced motion of the pαSyn after 14 weeks does not reflect a clear anxiogenic effect due to SNc DA cell loss, but most parsimoniously is explained by reduced locomotor activity. In conclusion, these results demonstrate that cell-type specific pαSyn-induced cell loss induces considerable locomotor impairments, with a timeline comparable to other AAV-driven rodent models^15^.

## DISCUSSION

We here use advanced analysis methods and readouts to characterize a new viral vector that delivers a floxed version of A53T-αSyn, thus enabling cell-type specific synucleinopathy via local injections into Cre-transgenic mouse driver lines. By specifically expressing pαSyn expression in DA-SNc neurons of DAT-Cre mice, we were able to recapitulate well-established features of synucleinopathy, namely i) pS129-positive pαSyn expression, a molecular marker for Lewy-bodies, within SNc neurons (*Fig. 2d*), ii) progressive degeneration of DA-SNc cells (*Fig. 2a-*c, *S3,4*), along with their nigrostriatal projections (*Fig. 2e-f*). This depletion is accompanied by iii) marked locomotor deficiency (*Fig. 3*). Notably, those cellular and behavioral effects arise gradually, but within a relatively short timeline (2.5 months) as compared to other genetic models of PD (14 months)^35^, providing compatibility with standard experimental approaches. Overall, our cell-type specific synucleinopathy model, by putting the focus on neuronal circuit dysfunction^36,37^, fills a gap in the existing toolbox for studying PD pathomechanisms.

Furthermore, there is increasing evidence for the importance of cellular heterogeneity as a major determinant of PD pathomechanisms. For instance, neuronal subtypes with distinct connectivity were identified in the SNc, suggesting different functions of neuronal subpopulations^36,37^. Among those roles, the relevance of neuronal subpopulations within a single region is documented by the fact that distinct optogenetic manipulation had different motor effects in PD mice^38^. In addition, differing vulnerabilities among DA neurons have been described^39^. We cannot exclude that our TH cell loss differences along the anterior-posterior axis are caused by varying susceptibilities that neurons show upon maladaptive αSyn presence. To investigate the progressive effects and vulnerability of αSyn aggregation within functionally defined circuit modules, these findings need to be refined by inducing a more specific pαSyn-driven circuit modification.

Methodologically, the versatility of the Cre-loxP system offers the possibility to use state-of-the-art methods for circuit targeting and manipulation^40^ in addition to progressive cell loss mimicking. For instance, a variety of other cell-type specific methods could be considered, such as i) optogenetics to modulate neuronal activity, ii) cell activity observation via calcium imaging or fiber photometry. Besides, our viral approach enables to choose unilateral alteration and keep the contralateral side as a control, or bilateral alteration as in putative PD.

A recurring challenge in characterizing neurodegenerative processes in PD has been to achieve reliable cell quantification^28,41^. New bioimage analysis approaches aim to overcome the disadvantages of manual cell counting^42–44^, which often depends on non-objectifiable principles and is at risk of observer bias. Machine learning-assisted models, which are generated from ground-truth assessed by multiple scorers perform better than networks trained by a single expert, as they pool different bases of knowledge^28^. Those automatized pipelines also speed up the process of cell counting, enabling high throughput of data. However, starting from input data conversion to a specific format, until postprocessing with reliable exclusion criteria and results validation represent numerous steps that are not implemented within the existing counting tools^27,29^. With *findmycells*, we developed an open source, deep learning-based end-to-end bioimage analysis software, usable by non-coding experts.

Mouse video-tracking on the rotarod unveiled finer observations of gait parameters. It revealed that not only the latency to fall is reduced, but also that mice are located lower on the rod. Interestingly, when inspecting mice behavior in the OF, we confirmed that DA-SNc targeting of pαSyn-driven degeneration induces locomotor symptoms, but without specifically affecting anxiety-like behavior. These results are in line with findings of enhanced anxiety before any DA-SNc loss^4,45^ and point towards the need for more specific circuit investigations to understand complex motor and non-motor symptoms in PD and the underlying circuitry. This is especially important considering that PD affects multiple monoaminergic network elements beyond the nigrostriatal DA system. Degeneration of neuromodulatory systems such as the noradrenergic^23,46^ and serotoninergic^47,48^ neuronal populations correlate with the appearance of non-motor symptoms^48,49^. By concomitant video- and heart rate recordings, our findings demonstrate that autonomic dynamics are affected by synucleinopathy. Further analyses are required to understand whether this is secondary to the locomotor impairments or due to direct effects on central autonomic circuits. Our strategy now allows to specifically investigate the role of individual neuromodulatory nuclei within the circuit at a given time, such as locus coeruleus^50^ and the dorsal raphe nuclei^51^, to address progressive loss of noradrenaline or serotonin, respectively. This enables refined pinpointing the spatial and temporal occurrence of the lack of those neuromodulators, the impacted targets, and their relationship with observed PD-like symptoms.

Through a more mechanistic understanding of PD pathomechanisms on the circuit level, our specific strategy could help unveil the contribution of individual pathways to existing treatments. Even if those proved their efficacy, they do not alleviate all symptoms^52^ and their precise mechanism of action remains unclear^3,53^. For instance, levodopa is hampered by side effects, like progressive on/off motor fluctuations or dyskinesia^3^. Further treatment options (e.g. deep-brain stimulation^54^) would also benefit from a detailed knowledge of basal ganglia network dynamics.

Nonetheless, some limitations should be considered. Because of the SNc’s extension along the anterior-posterior axis, potential heterogeneity of the injection site can cause variability of viral spread. Additionally, specific pαSyn expression does not resemble the kinetics of pαSyn toxicity occurring in multiple nuclei in parallel within the brains of PD patients^55^. However, while there is no animal model of PD that fully mimics all aspects of the disease^56^, our approach adds a complementary perspective, helping to further understand already established mechanisms with a focus on circuit-specific processes.

Our cell-type specific pαSyn targeting, thereby inducing targeted neurodegenerative processes, and associated behavioral impairments, analyzed with sophisticated impartial methodologies, constitutes the first step for detailed studies of networks effects of PD-like synucleinopathy. Targeted at a circuit level, our model opens a new scope of possibilities on unveiling the pathomechanisms of the disease.

## METHODS

### Animals

DAT-Cre (Slc6a3tm1(cre)Xz/J, stock n°020080) mice were obtained from Jackson laboratory and bred with 2- to 10-months old C57BL/6 wild type in an in-house animal facility. All mice were individually sheltered on a 12h/12h light-dark cycle. Experiments were carried out during the light cycle. Water and food were accessible *ad libitum*. This work follows the ARRIVE guidelines (Animal Research: Reporting of In Vivo Experiments)^57^. Local veterinary authorities, together with animal experimentation ethics committee approved the animal procedures (Regierung von Unterfranken: authorizations 2532-2-1356 and 2532-2-1067; IACUC-21-2018).

### Viruses

We used AAVs to deliver constructs of interest with genetical specificity. The pAAV-hSyn-DIO-mCherry (titer 4.56 e+14 gc/ml) was produced in an in-house viral production facility using the plasmid ordered from Addgene (#50459). The pAAV2/9-pCAG-FLEX-alpha-synuclein(A53T)-3*Myc-WPRE (mutated pαSyn, titer 3 e+12 gc/ml) and pAAV2/9-pCAG-FLEX-alpha-synuclein-3*Myc-WPRE (non-mutated αSyn, titer 3.94 e+12 gc/ml) were custom designed and produced by Vector Biolabs. The viral construct developed in the study will be deposited in Addgene for public availability before publication. AAV1/2-pCAG-FLEX-emptyvector-3*Myc-WPRE (empty vector, titer 5.16 e+12 gc/ml) was received from Chi Wang Ip’s team. To confirm the injection site targeting for cell loss assessment, the virus solution was mixed with blue beads (concentration 1:10). The AAV2/9-Cre^ON^-EGFP-KASH (titer 2.8 x 10E13 vg/ml) was custom designed and produced by VVF from the University of Zurich, and AAV2/9-Cre^ON^-H2B-Tomato (titer 9,09 e+12 gc/ml) was received from Silvia Arber.

### Stereotaxic surgeries

#### General procedure and virus injection

Male and female mice between 2 and 10 months-old were anesthetized with isofluorane (Harvard Apparatus, Iso-Vet Chanelle) mixed with O2-enriched air (induction: 4 %, maintenance: 1.5-2 %). For ensuring intraoperative analgesia, buprenorphine (0.05-0.1 mg/kg; Buprenovet, Bayer) was injected subcutaneously prior to the surgery. Throughout the surgery, body temperature was stabilized by using a far infrared warming pad (RightTemp Jr., Kent Scientific). When the breathing rhythm reached 1 breath/s, the mouse was taken out of the chamber and placed on the stereotaxic frame (Model 1900, Kopf). After ensuring total anesthesia by checking the pedal reflexes, local analgesia was administered by subcutaneous injection of ropivacaine under the scalp (8 mg/kg, Naropin, AstraZeneca). After midline incision and scull calibration, craniotomies were performed (0,4 mm diameter at 3.32 (±0.2) mm caudally, ±1.1 (±0.2) mm laterally from bregma, 4 (±0.2) mm depth to target the SNc. The virus solution was sucked into a glass capillary (calibrated micropipette 1-5 µl, Drummond Scientific) that was gently lowered to the target depth. 150nl of virus solution was injected at a velocity of 50 nl/min using a pressure injector (PDES-02DX, NPI electronic). The pipette was kept in place for 7-10 min for the virus to spread, before being slowly pulled out to avoid off-target expression of the virus. The scalp was further sutured and disinfected with antiseptics. To guarantee post-operative recovery analgesia, subcutaneous injection of meloxicam (5 mg/kg; Metacam, Boehringer Ingelheim) was made prior to suturing and every 12-24 h for 3 days. Finally, the mouse was placed back in a clean cage in front of a heating lamp to allow it to recover from the surgery. It was monitored and scored according to its state every day during 7 days after the surgery to ensure its good recovery.

#### Implantation of the electrode for an electrocardiographam (ECG)

To record cardiac parameters, ECG electrodes were implanted in all mice undergoing behavioral tests. ECG electrodes were custom built from micro connectors (A79108, Omnetics, MSA components) onto which three PFA-coated wires (7SS-1T, Science Products) were soldered. From those wires, two were dedicated to measure the ECG signal differentially, with the third wire being used as a reference. At the beginning of the surgery, two small incisions on the upper right and lower left torso were made following an imagined diagonal line centered on the heart. The two first wires were laced individually through a blunt feeding needle, placed subcutaneously from one skin incision of the ventral torso to the scalp cut. The wires’ tips were stripped from their insulation over 5 mm. A round-shaped glue tip was created and used to suture each wire onto some thoracic muscle tissue. The skin was then closed and treated with antiseptics. The connector was glued on the skull, with the third wire placed under the scalp.

### Histology

#### Perfusions and cutting

Mice were injected intraperitoneally with a Ketamine (100 mg/kg) and Xylazine (10 mg/kg) mix. After ensuring total anesthesia, they were transcardially perfused with first 1x PBS and then 4 % paraformaldehyde (PFA) for 5 min. Brains were postfixated in PFA 4 % at 4 °C overnight. For cell loss assessment (*Fig. 1&2*), brains were then washed in 1x PBS and incubated in 30 % sucrose for 72h before being cut in 30 µm-thick coronal slices using a cryostat (Leica CM1950). Since we observed that the viral expression was widespread, making it visible even in thicker slices, we used another protocol for brain processing following behavioral experiments (*Fig. 3*). In this case, the brains were washed with 1x PBS after PFA fixation, embedded in 6 % agarose cubes and, using a vibratome (Leica VT1200), cut in 60 µm-thick coronal sections. To reliably distinguish between the contralateral and ipsilateral sides, a sagittal cut throughout one side was made with a scalpel prior to cutting. For experiments in Fig. S4 brains were cut in 60 µm-thick coronal slices using a cryostat (Microm, HM 550).

#### Immunohistochemistry

Immunohistochemistry was performed in free floating sections. A first blocking of the slices was performed for 2 h at room temperature with 5 % Donkey Serum, 0,3 % Triton 100x in 1x TBS-T. The primary antibodies solution consisting in a mix of Myc-tag 1:500 (mouse, Ab32), TH-tag 1:500 (chicken, Ab76442 or sheep, Invitrogen PA1-4679), diluted into blocking solution. After 72 h of incubation at 4 °C, the slices were washed 3x 10 min in the blocking solution. The secondary antibodies solution was then applied for 2 h at room temperature. It contained a mix of donkey anti-mouse 1:1000 (Cy5, Jackson 715-175-150), donkey anti-chicken 1:800 (A488, Jackson 703-545-155) or donkey anti-sheep 1:800 (A488, Jackson, 713-545-147), donkey anti-rabbit 1:1000 (Cy3, Jackson 711-165-152) diluted in blocking solution. The slices were then incubated with DAPI solution (5 mg/ml 1:5000 in 1x TBS-T) for 5-10 min at room temperature. Finally, the slices were washed 2x 10 min in 1x TBS-T and 1x 10 min in 1x PBS. The slices were mounted on slides (SuperFrostPlus 25 x 75 x 1,0 mm, VWR 631-0108) and embedded in a custom-made glycerol-based mounting medium (Fluorostab) with 24×50 mm coverslips (Epredia). For Fig. S4 a similar protocol was used but the primary antibody mix consisted of mouse anti-GFP 1:1000 (Sigma G6539), rabbit anti-RFP 1:5000 (Rockland 600-401-379), and chicken anti-TH 1:500, (Neuromics CH23006) or chicken anti-Myc (Invitrogen A21281) diluted in blocking solution. The secondary antibody mix consisted of donkey anti-mouse 1:1000 (Alexa Fluor 488 ab150117, Abcam), donkey anti rabbit 1:1000 (Cy3, Jackson 711-165-152), and donkey anti-chicken 1:1000 (Cy5, Jackson 703-605-155) diluted in blocking solution.

A slightly different immunohistochemical protocol was used to stain Lewy bodies, based on the protocol by Henrich et al.^11^. Initially, the sections were washed 3 x 5 min in PBS and blocked for 2 h with 10 % donkey serum and 0,3 % Triton 100x in PBS. The sections were incubated with the primary antibodies rabbit anti-pS129 (1:2000, Abcam, ab51253) and goat anti-Myc (1:500, Abcam, ab9132) in PBS with 10% donkey serum for 1h at RT and then overnight at 4°C. The following day, sections were initially washed 4 x 5 min in PBS and incubated with the secondary antibody mix for 1 h (biotinylated anti-rabbit, 1:1000, Jackson, 711-065-152), then washed 3 x 5 min with PBS and finally incubated for 2 h with a fluorophore-conjugated streptavidin (Alexa Fluor 633, 1:1000, ThermoScientific, 21844) and donkey-anti-goat (Cy3, 1:800, Jackson 705-165-147) in PBS. Lastly, the sections were stained with DAPI and washed as described above. Sections were mounted using Polymount (Polysciences,18606-100).

### Imaging

#### Microscopy

Overview images from the brains were taken using a fluorescence microscope (Zeiss AxioImager 2 with a Plan-Apochromat 5x/0.16 M27 objective, or the Zeiss AxioZoom V16 equipped with an Apo Z 1.5x objective, magnification 50x). Images of TH-A488 in the SNc and striatum were taken with the AxioImager using the Plan-Apochromat 20x/0.8 M27 objective and 3×3 binning. We took z-stacks images using LED-Module 475nm (power 10 %, exposure time 0,75 ms) for DAPI and the LED-Module 488nm (power 40 %, exposure time 9 ms) for TH. Image acquisition was set using the Zeiss Tiles module. For cell loss assessment, the section with the blue beads was chosen as the center of viral injection. In addition to this section, the 3 sections anterior and posterior (every 60 µm) were imaged. If blue beads could not be clearly located in any section, the mouse was excluded from quantification. For the brains processed after behavior and Fig. S4, the slices were aligned to the Allen Brain Atlas and every 60 µm section ranging from 3.02 to 3.52 mm posterior to bregma was taken for the quantification.

Images of the pS129 staining and colocalization with pαSyn were done using a confocal microscope (Olympus IX81) with a 60x oil immersive objective (Olympus UPlan SAPO), together with a laser box (Coherent OBIS LX/LS) and a fluorescent lamp (Excelitas X-Cite 120Q). Images for Fig. S4 were taken with a fluorescent microscope with a 20 X objective (Zeiss Axio Observer 7) and a confocal microscope with a 10X objective (Zeiss LSM 980).

#### Image processing

For the analysis, the z-stacks were then converted in 2D images using either the maximal orthogonal projection or extended depth of focus functions of the Zen Blue Software, depending on the best signal outcome. ImageJ (https://imagej.nih.gov/ij/index.html) was used to define the regions of interest (ROIs) based on TH stainings in the Alexa 488 channel.

### Density analysis of the striatum

From single plane 5x images (6 per brain), the striatum borders (including caudate putamen and globus pallidus) were drawn according to the Allen Brain Atlas (https://mouse.brain-map.org/static/atlas). The mean density quantification tool from ImageJ was used on those ROIs.

### Quantification with *findmycells*

The methods implemented in *findmycells* are detailed in *Fig. S2*. In the post-processing step, identified features that were smaller than the experts’ labels, as well as those with no contact to a predefined region of interest (ROI) covering the SNc were excluded from the quantification.

### Behavior

Behavioral recordings were completed in sound-attenuated chambers (size: length: 100 cm, width: 80 cm, height: 116 cm) illuminated from the top by a circular LED lamp (LED240; Proxistar). The chamber used for open-field paradigm was covered with black insulating foam, illuminated at 350 Lux and contained a petri dish filled with 1 % acetic acid. The second chamber, used for rotarod recordings, was covered with white insulating foam, illuminated at 650 Lux and included a petri dish filled with ethanol. For each paradigm, the suitable context was placed in the center of the chamber and cleaned with the appropriate liquid before each recording (i.e. acetic acid/ethanol). The context temperature was kept at 22.5±1 °C. The paradigms were repeated at weeks 1, 4, 8, 12 and 14 after viral injection to assess cell loss progression on behavior.

#### Open field (OF)

For testing innate anxiety and naturalistic locomotion. The paradigm consisted in a 50 x 50 x 50 cm white box, in which animals were placed and left to freely explore for 15 min.

#### Accelerating Rotarod

For testing motor coordination, we used the accelerating Rotarod test (Ugo Basile). Our protocol was based on Esposito *et al*. 2014^58^. The trials consisted of 5 min-long recordings, during which the rod rotation velocity ramps from 5 to 50 rpm. Each trial was separated by a 5 min break. We started with a training phase one week before surgeries, in which animals underwent 5 trials per day for five days. During the experimental phase, to keep the mice habituated to the paradigm, they were trained twice, at 10 and 5 days prior to the testing day. On recording days, each mouse was tested 4 times. We always excluded the first trial, because of the intra-individual high variability we observed. We then averaged the latency to fall from the 3 last trials.

#### Recording system

The acquisition system (Plexon, Omniplex system) was recording analog and digital signals. It was combined with CinePlex Studio (Plexon) for synchronizing with top RGB camera videos (Pike Camera F-032C, Allied Vision, Campden Instruments). To control the experimental timings, the Radiant software was used through the PlexBright analog and digital outputs. An RZ6 multi-processor (Tucker-Davis Technologies) contributed to online processing and synchronization via a MATLAB/ActiveX/RPvdsEx interplay (MATLAB 2022b, The MathWorks; RPvdsEx, TuckerDavis Technologies). An initial train of TTLs followed by a 1Hz signal was generated and transmitted to the several systems for offline alignment. A custom GUI allowed pre-recording calibration, focus, and real-time visualization of the movie. After the signal was broadcasted by PlexBright and relayed to the RZ6, the recordings were triggered by MATLAB. They were similarly ended. The ECG signal was acquired with an amplifier (DPA-2FX, npi) linked to the OmniPlex system, at 5 kHz. According to the quality of the electrodes’ signal, it was recorded either differentially or from a single electrode.

### Heart rate analysis

Heart rate analysis was performed as described before^21^. In brief, the ECG signal was acquired with an amplifier (DPA-2FX, npi) linked to the OmniPlex system, at 5 kHz. Then, the analog signal was preprocessed, heart rate peaks were extracted, and heart rate was calculated as a sliding window within a custom MATLAB GUI (https://github.com/Defense-Circuits-Lab/ECGanalysis). Detection and retrieval of heart beats from the recorded ECG signals was done via a custom-coded MATLAB script. The raw signal was extracted from the .pl2 files, using Plexon’s SDK. If necessary, the frequencies were adjusted by bandpass filtering. The obtained signal was then elevated to the 4th power to boost separation and smoothed using a Gaussian filter. To extract the putative heart beats, a threshold was set. After obtaining the timestamps from this modified signal, heartbeat waveforms were withdrawn. The results were highly specifically pretreated using a divergent template-matching as well as interbeat interval confidence scoring algorithm. Ambivalent ranges were manually corrected by the experimenter. If that adjustment was impossible or in case of doubt by means of a bad signal-to-noise ratio (i.e. contamination by the electromyogram), the corresponding ranges were tagged and excluded from further analyses. Finally, the R peaks were extracted from each obtained waveform, and saved for additional processing.

### Open field analysis

Behavioral classification in the Open Field Test was performed as described before^21^. A custom-build MATLAB code was used to process the top view movies (https://github.com/JSignoretGenest/SignDCL/tree/master). For RGB movies, the 80th percentile of a selected number of frames was used to create a background image that was subtracted from all the frames. Then a threshold was manually chosen, and the obtained binary mask was adjusted following a succession of simple treatments: (i) morphological closing, (ii) removal of small objects, (iii) a second morphological closing, and lastly (iv) morphological opening. Parameters’ values were set to fit to each condition. Hence, for every frame, a mouse contour together with a center of gravity were generated from the resulting mask and finally saved. For later normalization, a calibration (movie pixels / cm size of real object ratio) was also generated by manually drawing a segment (usually the paradigm width) and indicating the corresponding length of the drawn object (in cm). In the next analysis steps, speed and mouse position were extracted from the coordinates of the center of gravity. They were also slightly smoothed via a median filter. Furthermore, to recapitulate general activity even when absence of locomotion, a motion measure was computed: it represents the percentage of pixel change within the mouse masks from one frame to the adjacent one (non-overlapping pixels/total pixel count).

#### Spatial analysis

From the contours drawn during the tracking step derived the OF areas. They were completed by manual delineation if needed. The OF was divided into a 5 x 5 grid. The center corresponds to the middle 2 x 2 square, the corners to the corners of the whole grid, and the corridors to the squares connecting two corners. Raw heat maps were obtained from the corresponding data and the tracking information with the FMA Toolbox (http://fmatoolbox.sourceforge.net/) using 250 bins without smoothing. A post-processing 2D smoothing was done ignoring the bins with no occupancy to avoid “edge” effects. The corresponding quantification was computed separately from heat maps, by extracting from the raw data the time bins for each area and averaging it. For the transitions between areas, they were considered as synchronizing events for the PSTH because of the lack of events matching this criterion.

### Rotarod analysis

The latency to fall was averaged across the 3 last trials and compared between weeks. For measuring gait parameters on the rod, mice were recorded from the back and DeepLabCut^33,34^ was used for markerless pose estimation. The resulting tracking data was processed using custom-written python code. The base of the tail was used as the position of the mouse on the rod (relative to the bottom left corner of the rod). Data was split and averaged across sessions, separated between a slow rotation phase, including all values when the rod spined between 6 and 10 rpm, and a fast rotation phase when the rod rotated between 10 and 50 rpm. Additionally, the heart rate and mouse position were observed when the mice were falling using peristimulus time histogram. For statistics, the heart rate was normalized to the mean heart rate from 20 s to 11 s before the fall and then the mean heart rates were compared between a bin from 11 s to 6 s and a bin from 5 s to 0 s before the fall. Similarly, the average position of the mouse on the rod was compared between 11 s to 6 s and 5 to 1 s prior to the fall.

### Statistics

For all cell count and density quantifications, the results were averaged and normalized to the highest value of the dataset. Normality was controlled with the Lilliefors test for each set of data, as well as homoscedasticity using the Brown-Forsythe test. When comparing only two sets of normally distributed data, a Student’s t test was applied, otherwise the Wilcoxon signed-rank test or the Mann Whitney U test were used. When comparing more than two series of data, if the hypotheses of equal variance and normality were true for all sets, a two-way ANOVA test was applied (or a mixed model in case of missing values), in both cases followed by corresponding post-hoc test with Bonferroni correction, otherwise Wilcoxon or Mann Whitney U tests were used for each comparison separately. A 3-way ANOVA was used for analyzing behavior x time interactions within OF results, using a custom-made MATLAB code together with Prism 9.1.5 (GraphPad). Interpretation followed a decision tree previously described^61^. The latency to fall scores from rotarod analysis were statistically evaluated using a linear regression, alongside individual repeated measures ANOVAs between weeks.

### Data and Code Availability

Data generated in this study are provided in the Source Data file. We provide a publicly available dataset of images and a *deepflash2* model for testing the functions of *findmycells*^62^. The code for *findmycells* is available on GitHub: https://github.com/Defense-Circuits-Lab/findmycells, alongside with an interactive documentation to easily install and use the tool: https://defense-circuits-lab.github.io/findmycells/tutorials/gui_tutorial.html.

## ACKNOWLEDGMENTS

We thank K. Walter, H. Troll and C.A. Mehling for technical assistance, M. Gruber for experimental support, as well as M. Moradi for support regarding confocal microscopy. We thank N. Wenger for valuable comments on our manuscript. We thank Silvia Arber for providing AAVs. Further, we thank all members of the Defense Circuits Lab for the discussions and support with the project. We also thank R. Sendtner and all members of the animal facility. We thank the Medical Informatics Service Center (SMI) of the University Hospital Würzburg for providing an infrastructure for data storage and analyses, and for technical assistance. This work was supported by the Deutsche Forschungsgemeinschaft (DFG, German Research Foundation) [Project-ID 424778381-TRR 295, B06], Heisenberg professorship to P.T.: [TO 1124/2], the Graduate School of Life Science, Medical Faculty of the University Hospital Wuerzburg (Fellowship to K.K., A.L., N.S.). S.R.R. is supported by the Add-On Fellowship of the Joachim Herz Foundation, Germany. C.W.I. received funding from the DFG [Project-ID 424778381-TRR 295, A06], the Interdisciplinary Center for Clinical Research (IZKF) at the University of Würzburg (S-506, N-362) and by the VERUM foundation. M.S.E. was supported by funding from NARSAD Young Investigator Grant by the Brain and Behavior Foundation, Career Development Award from HFSP, Neurodegeneration Challenge Network from the Chan Zuckerberg Initiative, CONICET and FONCYT (PICT-PRH-2019-00006, PICT-2019-00901) and IBRO regions connecting award.

## AUTHORS CONTRIBUTION

A.L., D.D., M.S., N.S., M.S.E and P.T. conceived the project and designed the experiments. A.L., N.S., S.R.R., K.K. and P.T. wrote the manuscript with contributions from all other authors. A.L., N.S., M.S.E., A.I.T., K.K. and S.R.R. performed histochemical experiments. A.L., M.S., A.I.T. and K.K. ran behavioral experiments. D.D., J.S.G, K.K., and M.S. set up and designed the analysis pipeline. A.L., D.D., J.S.G., K.K., M.S., N.S. and S.R.R. analyzed the data. M.S.E. provided the viruses and supported the experiment designing. C.W.I. provided the empty vector virus, as well as his expertise regarding the viral model approach and analyses.

## COMPETING INTERESTS

All authors declare no financial or non-financial competing interests.

## Notes

### Competing Interest Statement

The authors have declared no competing interest.

