## Supplementary figures for "A viral vector model for circuit-specific synucleinopathy"

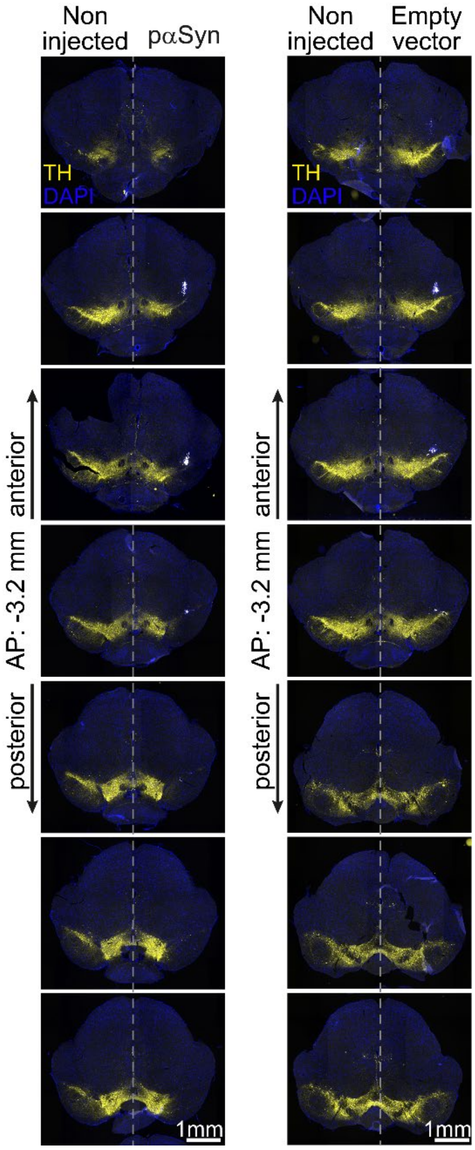


**Fig. S1 TH staining in SNc along the AP axis.** Representative images from anterior to posterior constituting a batch of image analysis, with the maximum blue beads signal on the 4th one.

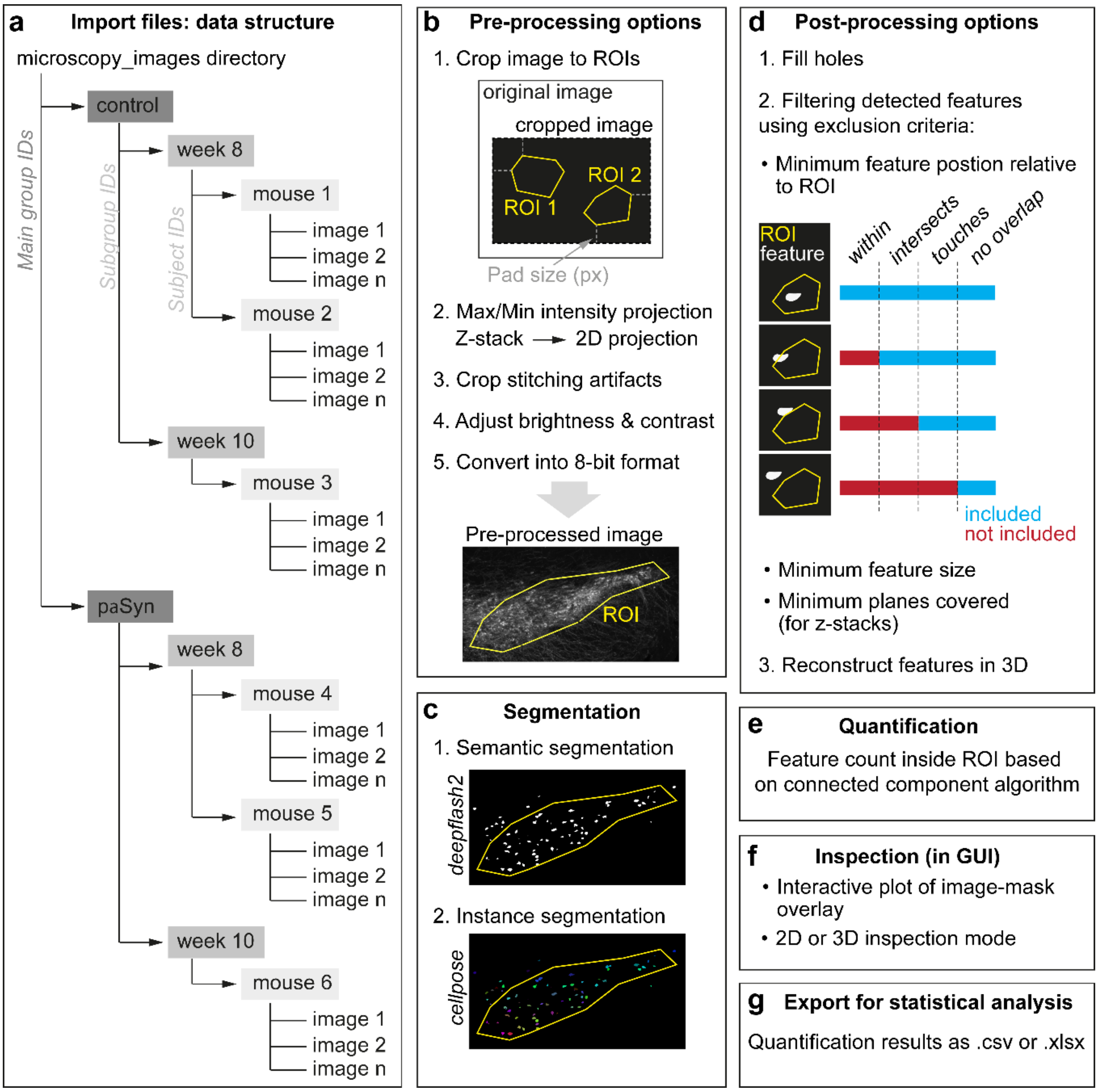


**Fig. S2 Findmycells pipeline in detail. (a)** Import. Image data in standard file formats (czi, .png, .tif) are supported. Using findmycells is possible with standard 2D as well as with z-stack images. To quantify cells in certain regions of the image, ROIs can be defined prior to the analysis, and loaded with the images. **(b)** Preprocessing. To reduce computational load in the segmentation process, any of the following preprocessing strategies can be applied to the images: 1) ROIs: if the analysis doesn't include the full image, but only one/some specific parts of it. In this case, it is possible to specify ROIs using ImageJ. Regions of the images that do not contain ROIs are cut off. The parameter "pad size" allows to keep some additional distance around the ROIs. Applying the strategy "crop image to bounding box enclosing all ROIs" will significantly reduce computation time, required memory and disk space for the later steps. 2) Z-stack projections: for z-stack images, to convert them to 2D, projection into one plane is possible by either using the maximum or the minimum pixel value from the individual image planes. The strategy "maximum intensity projection" uses the highest, the strategy "minimum intensity projection" uses the lowest pixel value. 3) Stitching: as microscopy images are stitched together from individual tiles, they might be slightly offset from each other. This can cause artifacts at the borders of the tiles in the form of usually fully black or fully white pixels. The strategy "crop stitching artifacts" aims at removing those pixels, as they can interfere with normalization of the images during segmentation strategies. 4) Brightness & contrast: can be automatically adjusted using the strategy "adjust brightness and contrast" by specifying the parameter "percentage of pixels that will be saturated". The strategy uses the function exposure.rescale_intensity from the scikit-image package[63]. The default value of saturated pixels is 0.35 %, which is also the standard value in ImageJ. For z-stack images, this strategy can be applied to the full z-stack or each plane individually via the parameter "adjust on whole z-stack level" if set to "globally", or on each plane individually if set to "individually". 5) Bit depth: depending on microscopy settings, images can be acquired at different bit depths. Deepflash2 and cellpose, the two packages used for segmentation, require 8-bit images as input. Therefore, as the last preprocessing strategy "convert into 8-bit format" must be applied. **(c)** Segmentation. For segmentation of the images, a deepflash2 model ensemble[27] must be trained on the specific dataset prior to the findmycells analysis. Selection of the training data is a crucial part, since deepflash2 can only predict features in experimental images reliably, if they are similar to the training images. We refer to the deepflash2 documentation[64] for a detailed explanation of how to train a new model. Applying deepflash2, provides a first categorization of the image pixels into cell or no-cell, so called semantic segmentation. A likelihood threshold of 0.5 was applied to the resulting segmentations. In a second step, cellpose[29] can be applied to get instance segmentations. Cellpose offers publicly available generalist models trained on a variety of images and can be used without the need of retraining in findmycells to predict nuclei or cells (nuclei, cell models). Further information on these models can be found in the documentation of cellpose[65,66]. Recommended for this step, especially on large datasets, is using a graphical processing unit (GPU). We provide a Google Colab notebook and the documentation[67] for installing the required packages and using a GPU. **(d)** Postprocessing. Any of the following postprocessing strategies can be applied to the images depending on which detected feature you want to keep for quantification: 1) Fill holes: during segmentation, holes inside of labeled features can be detected. Under some conditions, those gaps can represent the truth. That's the case if the cell's cytoplasm is detected sparing out the nucleus, or if imaging biconvex erythrocytes in one plane. Based on the research question, for instance if the wished outcome is the total cell counts, the gaps can be considered artificial. The holes are removed by application of the strategy "fill holes", which uses the function ndimage.binary_fill_holes from the scipy package[68]. 2) Exclusion criteria: to filter the detected features and include only certain categories into quantification, it may be necessary to apply certain exclusion criteria. This can be done using the strategy "filter detected features using exclusion criteria". Features categories depend on their position according to the ROI. Features lying completely inside the ROI borders are classified as "within". If the features are intersected by the area ROI border, they are named "intersects". The classification "touches" describes features in contact with the border of a ROI. Every feature completely outside the ROI borders is labeled as "no overlap". All features that meet the classification specified as "minimum feature position relative to area ROI", or being even closer to the ROI, will be included. In z-stack images, a feature's position can be classified differently in different image planes. In this case, the feature will be in- or excluded depending on the nearest classification. The feature size can be used to distinguish between valid and artificial features. Providing a "minimum feature size in px" will exclude any feature with a smaller area. If this strategy is applied to a z-stack image, the feature size is determined in the plane with its largest expansion. For z-stack images, the number of consecutive planes that each valid feature must cover can be defined. If a feature is present in fewer planes than the number of "minimum planes covered", it will be excluded. 3. 3D: features detected in 2D images belonging to z-stack images, can be merged by findmycells using the strategy "reconstruct 3D representations of cells". Features are considered as coherent, if there is an adequate overlap across planes. If multiple features overlap, only the one with the largest overlap is considered as overlapping. **(e)** Quantification. The number of unique features inside each ROI is determined. This step is based on the application of an implementation of the connected-component algorithm in python, referred to as cc3d69. **(f)** Inspection. To allow users to reconstruct how the images were changed in the individual steps, the images are saved to disk after each step. Apart from that, it is possible to create interactive plots within findmycells that allow for a closer look on detected 2D and 3D reconstructed features. **(g)** Export. Detailed tables with the quantified results can be exported in .csv or .xlsx format for performing statistical analysis.

To complete the explanations, a step-by-step tutorial is available at https://defense-circuits-lab.github.io/findmycells/tutorials/gui_tutorial.html

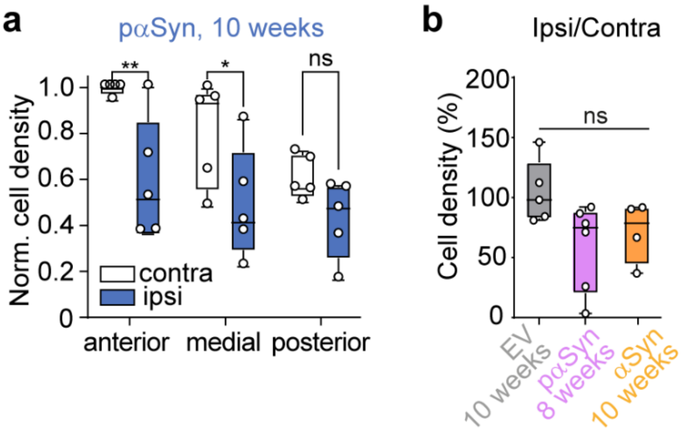


**Fig. S3 Cell quantification following paSyn expression (8- and 10-weeks post-injection) and non-mutated αSyn expression. (a)** Quantification of TH+ cell density along the antero-posterior axis in the ipsi- and contralateral SNc of mice injected with pαSyn (10 weeks post-injection). Box plots show mean ± min/max values, n = 5 mice, two-way ANOVA with Sidak's multiple comparisons test (*p<0.05). **(b)** Percentage of ipsilateral/contralateral cell density for EV (n = 5 mice; same data as Fig. 2c), pαSyn (8 weeks post-injection, n = 6 mice) and non-mutated αSyn (10 weeks post-injection, n = 5 mice). Dots correspond to mean values per animal. Box plots show mean ± min/max values, Mann-Whitney test (*p<0.05).

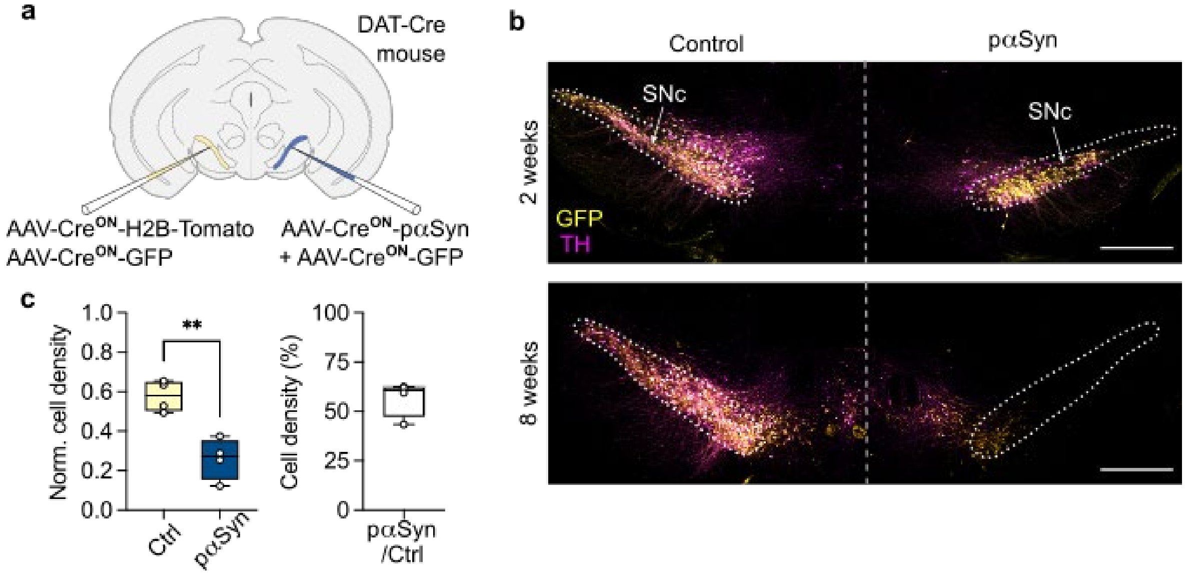


**Fig. S4 Viral tagging approach to quantification of neurodegeneration effects. (a)** Summary of experimental design. **(b)** Representative maximum-intensity projections from confocal z-stack images showing expression of GFP (yellow) and TH (magenta) in SNc neurons after 2 or 8 weeks of virus injection. Note the reduction in GFP+ and TH+ labeling on the side injected with pαSyn. Scale bar = 500 μm. **(c)** Quantification of GFP+ cell density in the left (control, Ctrl) and right (injected with pαSyn) SNc (left panel), and percentage of right/left GFP+ cell density in SNc (right panel) after 8 weeks of expression (n=4). Circles correspond to mean values per animal. Paired two-tailed t-test. (**p<0.01)

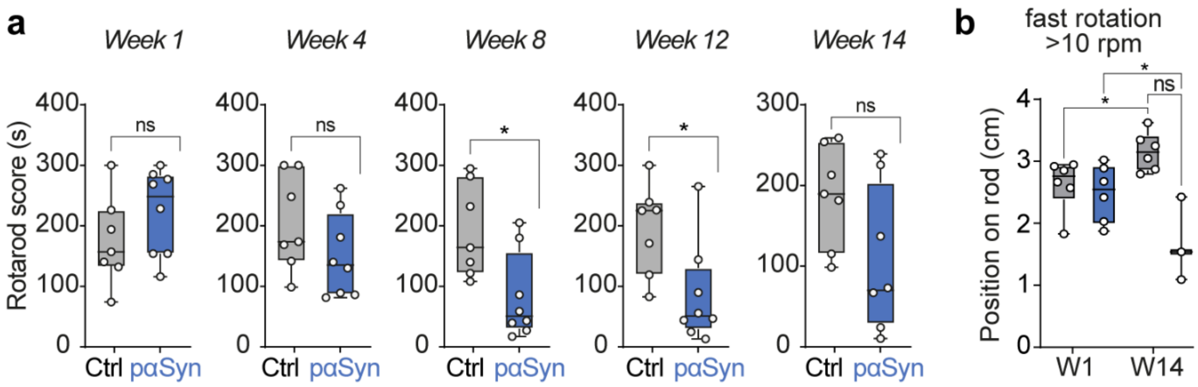


**Fig. S5 Latency to fall on the rotarod. (a)** Rotarod scores for pαSyn and control mice at different time points after AAV injection (weeks 1, 4, 8, 12 and 14). Dots correspond to individual values, n = 8 and 7 mice for pαSyn and control, respectively. Box plots show mean ± min/max values. Mann-Whitney test. **(b)** Mean tail base position during rotarod trials at fast rotation (10-50 rpm) at weeks 1 and 14. Box plots show mean ± min/max values, n=6 mice for both pαSyn and control. Mixed model with Bonferroni test (*p<0.05).

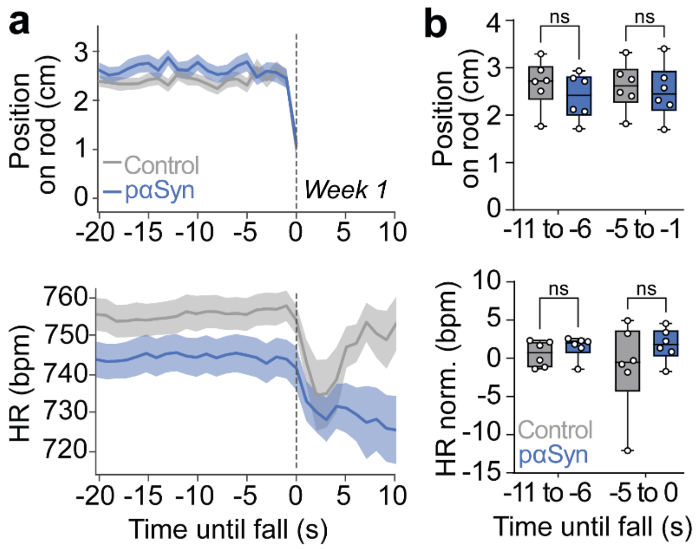


**Fig. S6 Quantification of rod position and heart rate on the rotarod at week 1. (a**) Peristimulus time histogram (PSTH) of the position of the mouse on the rod (top) and heart rate (bottom) from 20 s before, until 10 s after the fall. Dashed line shows when the fall occurs. **(b)** Comparison of mean position on the rod (top) and heart rate (bottom) for two timepoints. Mixed model with Bonferroni test.
